# Multi-area recordings and optogenetics in the awake, behaving marmoset

**DOI:** 10.1101/2021.10.30.466578

**Authors:** Patrick Jendritza, Frederike J. Klein, Pascal Fries

## Abstract

The common marmoset has emerged as a key primate model in neuroscience. Marmosets are small in size, show great potential as transgenic models and exhibit complex behaviors. These advantages place the marmoset model in the critical gap between rodents and larger primates. Thus, it is necessary to develop technology that enables monitoring and manipulation of the neural circuits underlying the behavior of the marmoset. Here, we present a novel approach to record and optogenetically manipulate neural activity in the awake, behaving marmoset. Our design utilizes a light-weight, 3D printed titanium chamber that can house several high-density silicon probes for semi-chronic recordings, while enabling simultaneous optogenetic stimulation. Surgical procedures are streamlined via custom 3D printed guides and implantation holders. We demonstrate the application of our method by recording multi- and single-unit data from areas V1 and V6 with 192 channels simultaneously, and show for the first time that optogenetic activation of excitatory neurons in area V6 can influence behavior in a detection task. Together, the work presented here will support future studies investigating the neural basis of perception and behavior in the marmoset.

## Introduction

The common marmoset (Callithrix jacchus) is becoming an important animal model in neuroscience (Solomon and Rosa, 2014; Miller, 2017; Servick, 2018; Okano, 2021). Due to its small size, genetic tractability (Sasaki et al., 2009; Tomioka et al., 2017; Sato et al., 2020) and complex behavioral repertoire (Stevenson and Poole, 1976; Mitchell and Leopold, 2015; Miller et al., 2016), it is placed in the critical gap between rodent models and larger primate models. Thus, marmosets hold great potential for improving our understanding of the neural circuits underlying complex behaviors and perception. It is therefore pivotal to develop techniques that enable monitoring and manipulation of these circuits in awake, behaving animals.

Many important technical advancements in neuroscience research with marmosets have been achieved in recent years. For example, the method of calcium imaging has been established as a promising optical alternative to monitor activity of individual neurons (Yamada et al., 2016; Kondo et al., 2018; Mehta et al., 2019). Nevertheless, extracellular single unit recordings remain the essential method in systems neuroscience due to their unparalleled temporal resolution and ability to record from almost any location in the brain (Steinmetz et al., 2018). Technical improvements in extracellular single unit recordings in awake marmosets were initially driven by the field of auditory research (Eliades and Wang, 2008; Remington et al., 2012; Roy and Wang, 2012). These recordings mostly utilized tungsten microelectrodes, which have limitations in terms of electrode density and geometric arrangement of recording sites. To overcome these issues, silicon-based microelectrode arrays have recently been established in awake marmosets (Johnston et al., 2019; Pomberger and Hage, 2019; Davis et al., 2020; Walker et al., 2021). These contributions have paved the way for better access to the neural circuits of the marmoset brain.

There exists a substantial body of work on the visual cortex of the marmoset (for comprehensive reviews, see Solomon and Rosa, 2014 and Mitchell and Leopold, 2015). However, the characterization of response properties of neurons is almost entirely based on experiments performed under anesthesia. In contrast, data from visual areas in awake marmosets is still very scarce (Porada et al., 2000; Johnston et al., 2019; Davis et al., 2020). Even more strikingly, there is only one study of single unit recordings in awake marmoset primary visual cortex (V1) (Porada et al., 2000), in stark contrast to the wealth of studies on this area in other species. Hence, the relative lack of published work in awake animals emphasized the need to develop suitable recording approaches.

Technical as well as conceptual advancements have revealed that computations in the brain are carried out by populations of neurons (Saxena and Cunningham, 2019). These populations are distributed within and across areas (Poggio, 2011; Panzeri et al., 2015). Thus, it is of great interest to be able to record from both, local populations, and from distributed populations across multiple areas simultaneously. For this reason, implant designs should be compatible with modern electrode technology, such as high-density silicon probes optimized for these applications, and they should ideally allow to target multiple brain regions simultaneously (Shobe et al., 2015; Steinmetz et al., 2021).

Importantly, beyond the correlative evidence that can be obtained from neural recordings, direct manipulation of neural activity can be used to gain insight into the causal link between neural circuits and behavior (Wolff and Ölveczky, 2018). Optogenetics is a powerful tool for such questions, because it offers the necessary spatiotemporal and genetic precision (Fenno et al., 2011). The principal feasibility of optogenetic stimulation techniques in marmosets has already been demonstrated (Macdougall et al., 2016; Komatsu et al., 2017; Ebina et al., 2019). However, the integration of neural recordings, optogenetics and behavioral manipulation is still lacking. Therefore, the aim of this work was to integrate these components into a well-engineered design that enables state-of-the art experimental access in the awake, behaving marmoset.

The approach presented here is based on semi-chronic recordings from multiple high-density silicon probes. It makes use of a light-weight titanium chamber, fabricated with metal 3D-printing technology, while surgical procedures are streamlined by means of 3D printed guides and implantation holders. We demonstrate multi- and single-unit recordings from two visual areas with 192 channels simultaneously and show that optogenetic stimulation of visual area V6 can influence the animal’s behavior in a perceptual detection task. Thus, we demonstrate for the first time neural recordings and optogenetic stimulation in combination with behavioral manipulation in the awake behaving marmoset.

## Results

### Implant design and recording approach

Our goal was to design a small and lightweight implant that utilizes modern high-density silicon probes while providing access to optogenetic stimulation techniques in awake behaving marmosets.

The complete implant consists of multiple parts: Headpost, chamber, microdrives, stabilizers, silicon probes and printed circuit boards (PCBs) holding the connectors (Fig. 1a). The 3D printed titanium chamber was designed to smoothly fit onto the surface of the marmoset skull (Fig. 1a, b). This was achieved by using a computed tomography (CT)-based skull model as anatomical reference for the curvature of the bottom of the chamber. The chamber houses six PCBs with connectors, which relay the neural signals from two silicon probe arrays: A four-shank 4×32 channel silicon probe is attached to a microdrive targeting visual area V6. A two-shank 2×32 channel silicon probe is located at the posterior end of the chamber to target visual area V1, amounting to a total of 192 channels. Both probes are implanted in the left hemisphere (Fig. 1a).

**Figure 1.**
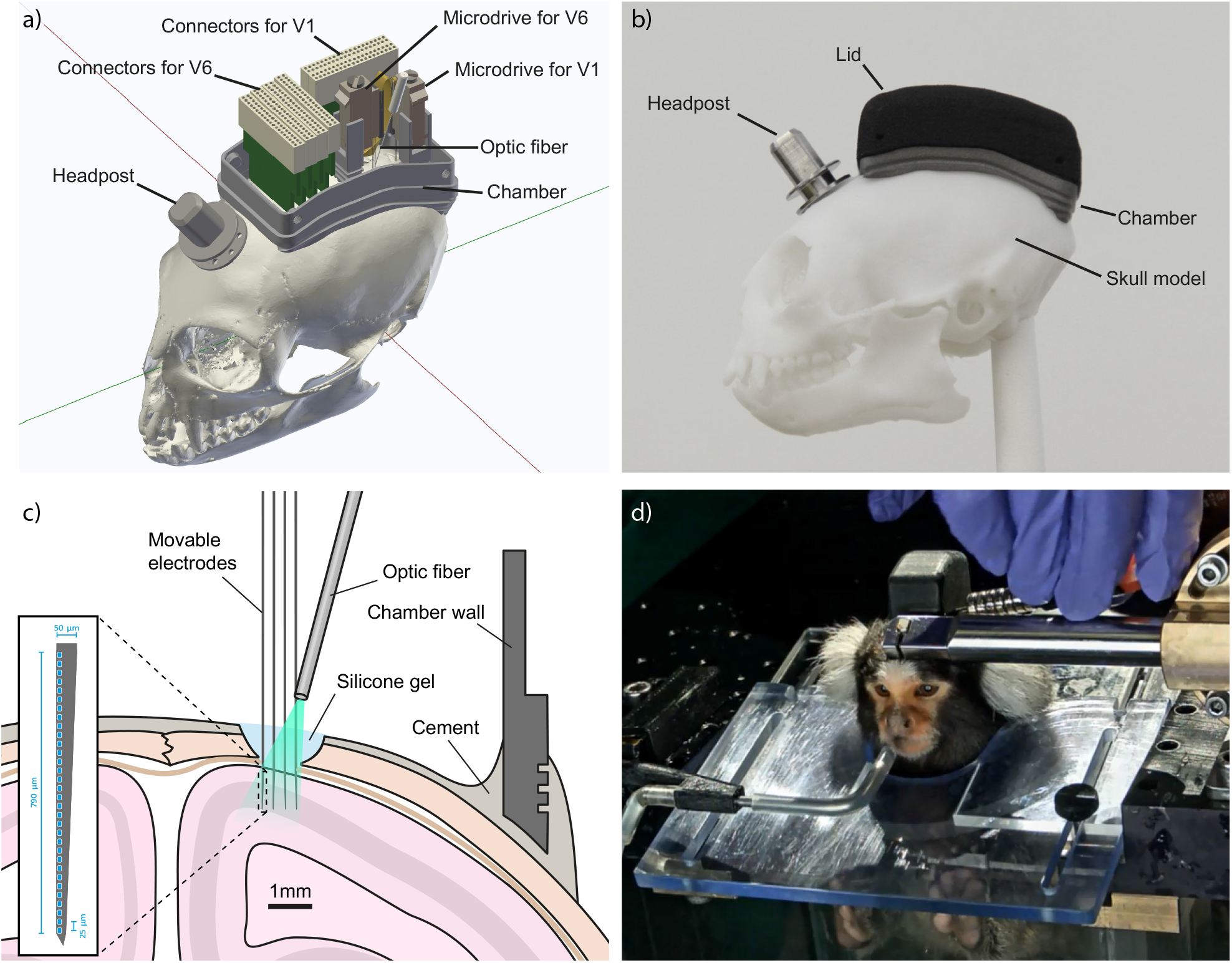
Implant design and recording approach. a) 3D rendering of the complete 192-channel implant design. A four-shank silicon probe with 4×32 channels is attached to the microdrive targeting area V6. A two-shank silicon probe with 2×32 channels is targeting area V1. The four connectors at the anterior end of the chamber are wired to the probe in area V6. The two connectors at the posterior right side of the chamber are routed to the probe in V1. An optic fiber (200 µm diameter) is placed above the V6 craniotomy with an external micromanipulator (not shown for clarity). The headpost for stabilizing the animal during recording is placed in front of the chamber. b) Side view photograph of a skull model with headpost, chamber and flat lid as used after implantation of headpost and chamber. c) Illustration of a coronal section of the target location in area V6. Craniotomy, electrodes and chamber are drawn to scale. Inset shows magnified view of electrode layout. d) Photograph of Monkey A while head-fixed and facing the monitor, during opening of the tall lid used after electrode implantation. The photograph shows the animal with the final 192-channel implant.

It is often advantageous to be able to move electrodes to a new recording position after signal decay, or in order to target a particular depth within the brain structure of interest. Therefore, we mounted the probes to microdrives which allow for up to 5 mm vertical travel. This makes it possible to change the recording position along the depth axis if required. Both microdrives are attached to titanium stabilizers that are 3D printed from the same material as the chamber. The stabilizers are intended to provide additional rigidity after implantation. Furthermore, they minimize the gap between the bottom of the microdrive and the skull, which needs to be filled with cement during implantation. Thus, the stabilizers also make the implantation process easier and faster.

Silicon probes are implanted through a small (≈2 mm diameter) craniotomy (Fig. 1c). After superficial insertion of the probes into the brain, the craniotomy is sealed with a transparent silicone gel (Jackson and Muthuswamy, 2008). Optogenetic stimulation can then be performed by pointing an optic fiber at the craniotomy such that the light penetrates through the silicone into the tissue (Fig. 1c). The optic fiber is held by an external micromanipulator that guarantees flexible and precise positioning.

To allow stabilization of the animal’s head during recordings, a headpost was implanted in front of the chamber (Fig. 1a, b). Both, the headpost as well as its holder (Fig. 1d) were produced by standard CNC milling from medical-grade titanium (Ti6Al4V). 3D printing was not viable here, because it does not offer the precision necessary for the fit between headpost and its holder, without substantial post-processing (Chen et al., 2017). However, alternative headpost designs could overcome this limitation (see Discussion).

The inside of the chamber is protected by a 3D printed nylon lid that can be secured by four small screws on the side of the implant (Fig. 1a, d). Threads for the screws were manually added after 3D printing. The use of 3D printed lids makes it possible to rapidly and flexibly produce multiple versions of lids. Before electrode implantation, the inside of the chamber does not contain any parts other than the (optional) reference wires. Therefore, the initial version of the lid was flat and could later be replaced by a taller version. This procedure allowed the animals to gradually get habituated to the size and weight of the final implant. Fig. 1b, shows a photograph of the chamber on a skull model with the flat version of the lid and the headpost in place.

We implanted chamber and headpost in five animals (Table 1). All animals tolerated the implant well, without the necessity of post-implantation wound care. None of the 3D printed nylon lids did require replacement, even after several months of use with almost daily opening and closing. Three of the five animals were subsequently implanted with electrodes in areas V1 and V6. Figure 1d shows a photograph of the final implant in Monkey A during opening of the lid just prior to electrophysiological recording.

**Table 1:** List of all animals, procedures and outcomes:

|  | Monkey A | Monkey U | Monkey D | Monkey E | Monkey P |
| --- | --- | --- | --- | --- | --- |
| Sex | male | male | male | male | male |
| First surgery performed (head-post, chamber, ref. wire) | yes | yes | yes | yes | yes |
| Body weight at first surgery | 385g | 438g | 455g | 530g | 428g |
| Second surgery performed (electrodes, viral vector) | yes | yes | yes | no | no |
| Body weight at second surgery | 371g | 460g | 445g | n.a. | n.a. |
| Neural recordings in V1 | yes | yes | yes | n.a. | n.a. |
| Neural recordings in V6 | yes | poor | yes | n.a. | n.a. |
| Optogenetic stimulation in V6 | yes | poor | yes | n.a. | n.a. |
| Duration (months) after first surgery* | 40 | 26 | 26 | 26 | 26 |
| Duration (months) after second surgery* | 35 | 19 | 19 | n.a. | n.a. |
| Data shown in figures | Fig.1, Fig.4, Fig.5, Fig.6, Fig.7, Fig.8 | - | Fig.3, Fig.7 | - | - |

\*Relative to the time this manuscript was prepared (September 2021)

Size and weight minimization of an implant are important design factors when working with small animals. These factors are not only crucial in order to ensure the welfare of the animal, but also facilitate the study of natural behaviors (Kondo et al., 2018; Courellis et al., 2019).

The chamber was designed to span 28 mm in the anterior-posterior axis and 17 mm in the medio-lateral axis of the skull (Fig. 2 a, b; outer chamber dimensions). We restricted the lateral extent of the chamber such its implantation required only minimal detachment of the temporal muscle from the bone. Consequently, no resection of the muscle was necessary. The sides of the chamber extended laterally only 1-2 mm beyond the superior temporal lines of the skull. This design allows targeting a large number of dorsal brain areas for neural recording and stimulation (Suppl. Fig. 1).

**Figure 2.**
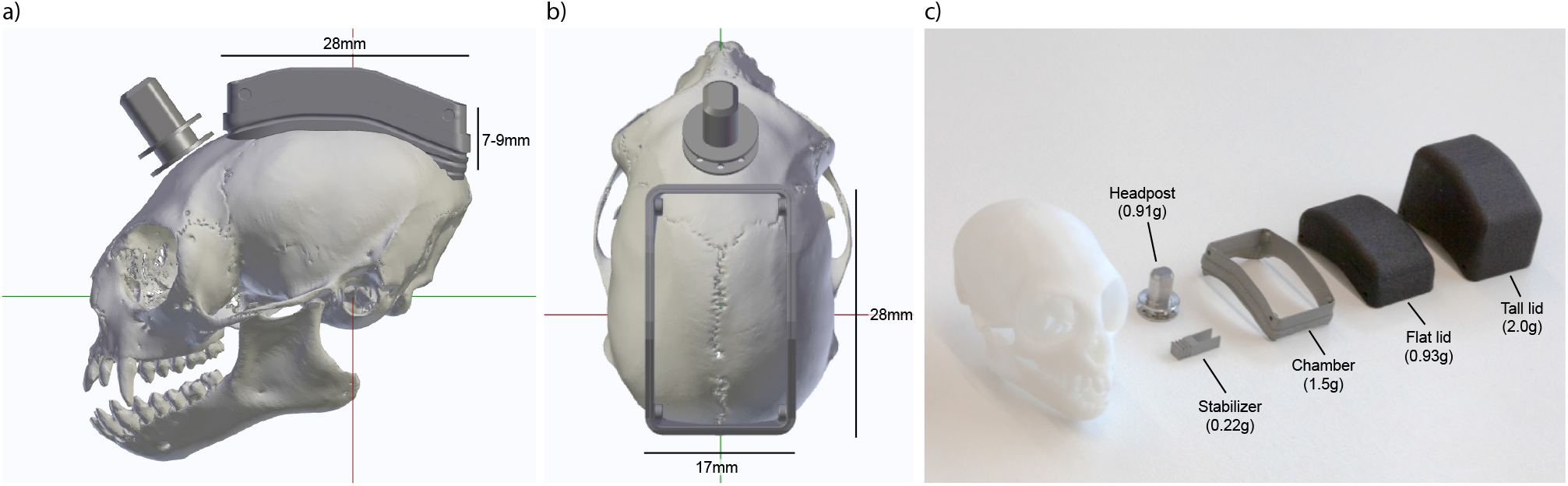
Implant size and weight. 3D rendering of side view (a) and top view (b) of a marmoset skull with headpost and chamber in target position, aligned in stereotaxic coordinates. Red line indicates interaural axis. Green line indicates anterior-posterior axis. c) Photograph of the CNC machined and 3D printed parts of the implant next to a skull model. Weights are indicated in parentheses.

The height of the final implant depends on the selection of electrodes and connectors inside the chamber. The chamber itself (without lid) protrudes only 7-9 mm from the surface of the skull. When closed with the flat lid (e.g. without probes installed), it reaches a height of 12-14 mm from the skull. After implantation with silicon probes and connector PCBs as used here, the chamber is closed with a taller version of the lid, and the implant reaches a height of 20-22 mm from the skull.

The total weight of the implant depends on its size and the density of the materials that are used. Recent advancements in metal 3D printing make it possible to accurately produce complex shapes from medical-grade titanium (Ti6Al4V). The mechanical strength of titanium allowed us to reduce the wall thickness of the chamber to 0.5-1 mm (Fig. 1c and Fig. 2b), which resulted in a weight of only 1.5 g for the chamber (Fig. 2c). Headpost and stabilizers had a weight of 0.91 g and 0.22 g, respectively. Lids were produced from a polyamide (PA12 nylon). Polyamides such as nylon show exceptional tensile strength, resistance to abrasion and can be 3D printed in a cost-effective way (O’Connor et al., 2018). Weights of the lids for the flat and tall version were 0.93 g and 2.0 g, respectively. Thus, the total resulting weight of the implant was approximately only 8 g, including headpost, chamber, silicon probes, microdrives, stabilizers, connectors and cement.

The implant design presented in this work combines several significant improvements over existing methods. It is small and extremely lightweight and enables recordings with a large number of channels as well as access for optogenetic stimulation. Because most parts are 3D printed, they can be manufactured very quickly at low cost and can be rapidly adapted for other methods such as calcium imaging or functional ultrasound imaging.

### Two-stage implantation procedure

Surgeries for experiments of the type described here often include a number of critical steps, such as: precise alignment of several independent parts, insertion of electrodes in multiple target areas and injection of viral vectors. Performing any of these steps is challenging even individually, and combining all of them in one surgery increases the risk of failure. To maximize chances of surgical success, we adopted a two-stage implantation procedure and made use of customized 3D printed implantation holders. First, headpost and chamber were implanted in the same initial surgery (Surgery 1). After appropriate recovery time, a second surgery was performed, in which a viral vector was injected and several silicon probes were implanted (Surgery 2).

### Surgery 1: Implantation of chamber and headpost

At the beginning of the first surgery, the animal was placed in a stereotaxic apparatus, and the skull was prepared for the implant (see Materials and Methods). Chamber and headpost could then be lowered onto the skull surface for alignment. Precise alignment of the chamber relative to the skull was crucial, because it ensured that the chamber could later be used as positional reference for the stereotaxic coordinate system. Both, chamber and headpost were held by a custom implantation holder that was attached to a micromanipulator (Fig. 3a). Prior to the surgery, cross-shaped markers on the sides of the holder were used for alignment to the interaural line (i.e. the axis of the ear bars). This assured correct positioning of the chamber in the anterior-posterior axis. During the surgery, a downward-pointing wedge integrated into the holder was aligned to the central skull suture, to assure correct positioning in the medio-lateral axis (Fig. 3a). After alignment, the position of the holder was locked, and the holder was temporarily removed to allow better access for the subsequent surgical steps.

**Figure 3.**
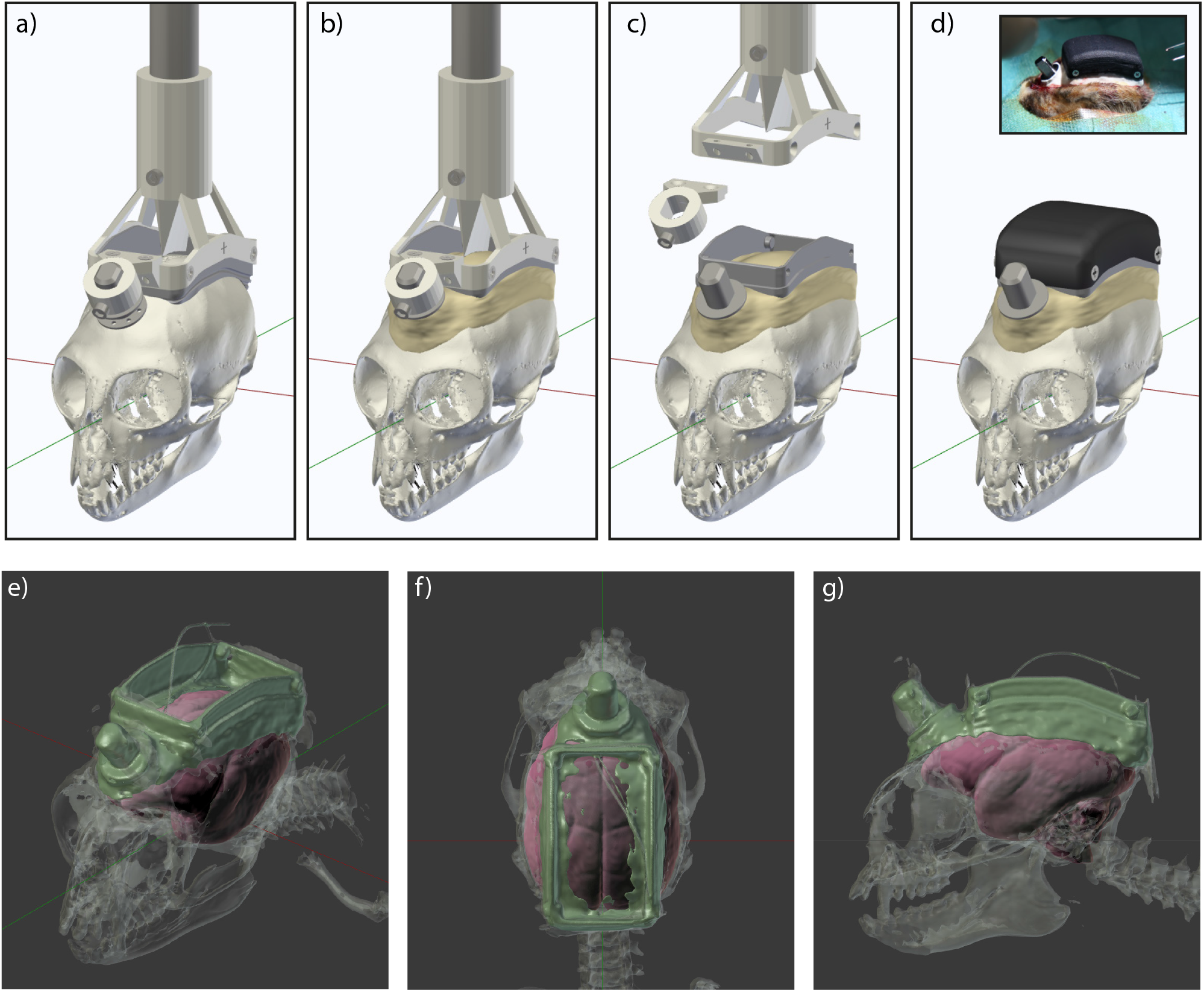
Surgery 1: Implantation of chamber and headpost. a) Chamber and headpost were held by a custom implantation holder that was attached to a micromanipulator. Note the cross-shaped markers on the side of the holder, used for alignment to the interaural axis, prior to the surgery. A wedge-shaped guide pointing downwards in the center of the holder was used for medio-lateral alignment to the central skull suture. b) Following skull preparation, the aligned chamber and headpost were cemented onto the skull. c) Once the cement had hardened, chamber and headpost were released from the holder. d) The chamber was closed with a 3D printed nylon lid for protection. Inset shows photograph of the implant at the end of the first surgery. e) Near-isometric projection, f) top view and g) side view of the 3D segmentation from a CT scan after the first surgery in monkey D. Radio-opaque cement, metal parts and reference wires show the highest contrast and are colored in green. Bone is shown in semi-transparent gray. The fitted MRI-based template brain is shown in red.

Marmosets have thin skulls and a narrow subdural space, which can make the use of bone screws problematic. Therefore, we used only dental adhesive and cement to secure the implant to the skull (Johnston et al., 2018). To this end, the skull surface was cleaned, roughened with a metal brush and coated with dental adhesive before a thin layer of cement was applied. Two platinum wires were implanted epidurally anterior to the chamber, serving as backup reference wires. Next, the implantation holder was returned to the previously determined antero-posterior and medio-lateral position, and lowered until the chamber contacted the skull. Following a final visual inspection of alignment, the headpost and chamber were cemented in place (Fig. 3b). After the cement had hardened, headpost and chamber were released from the holder (Fig. 3c). At the end of the surgery, the flat version of the 3D printed nylon lid was used to close the chamber (Fig. 3d). The animal was then allowed to recover for two weeks and subsequently underwent head-fixation training.

Variability in head morphology between animals can lead to inaccuracies during stereotaxic surgeries. Therefore, after the first surgery, we obtained anatomical data of the skull and implant via computed tomography (CT) scans (Fig. 3e-g). Appropriate thresholding of the CT images allowed segmentation of the bone (shown in transparent gray), and of metal and radio-opaque cement (shown in green). The cement layer in the center of the chamber was very thin and is therefore not visible everywhere in the segmented data, even though the skull inside the chamber was completely covered with cement. Also, note that the platinum wires appear thicker than they actually are due to the high CT contrast of the metal.

After segmentation, the inside of the animal-specific skull model was used to fit an MRI-based template marmoset brain (Liu et al., 2018). This approach can be justified under the assumption that the gap between bone and the brain is very small. Fits were performed manually by translating and scaling in all three spatial dimensions, and rotating in the pitch axis. The resulting fit of the template brain and its area delineations can then serve as individualized anatomical reference for each animal. Thereby, we obtained the precise positions of our target areas in the same reference frame as the chamber visible in the CT. Note that this CT-based targeting refinement was only used in marmosets D and U.

### Surgery 2: Injection of the viral vector and implantation of silicon probes

To assure correct positioning in the second surgery, the implantation holder from the first surgery (Fig. 3a-c) was used to re-align the animal’s head via the previously implanted chamber: After ensuring sufficient depth of anesthesia, the lid was removed, and the chamber attached to the animals’ skull was re-inserted into the holder. This effectively re-aligned the skull of the animal to precise stereotaxic coordinates as defined by the holder and the chamber. Subsequently, a high-precision articulated arm was used to fix the animals’ head position via the implanted headpost. After locking the articulated arm, the chamber holder was removed. Thus, the use of ear bars and eye bars could be avoided in the second surgery, thereby reducing potential discomfort for the animal.

Next, the inside of the chamber was disinfected with H_2_O_2_ and ethanol. A 3D printed guide was temporarily placed on the chamber and used to mark the target positions for the craniotomies over areas V1 and V6 of the left hemisphere (Supplementary Fig. 2). In Monkey A, coordinates for the guide were based on Paxinos et al., 2012, in monkeys D and U, coordinates were based on area delineations of Liu et al., 2018 after CT-based fitting to the individual animal, as described above.

Two platinum wires, serving as reference electrodes, were then implanted subdurally at the anterior end inside the chamber, through a small burr hole (≈2 mm diameter). Next, two small burr holes were made at the target locations for the electrodes over V1 and V6. A durotomy of approximately 1.5 mm was performed over area V6, and the viral vector was injected (Fig. 4a). After a short waiting time for diffusion of the vector into the tissue, the needle was slowly retracted.

**Figure 4.**
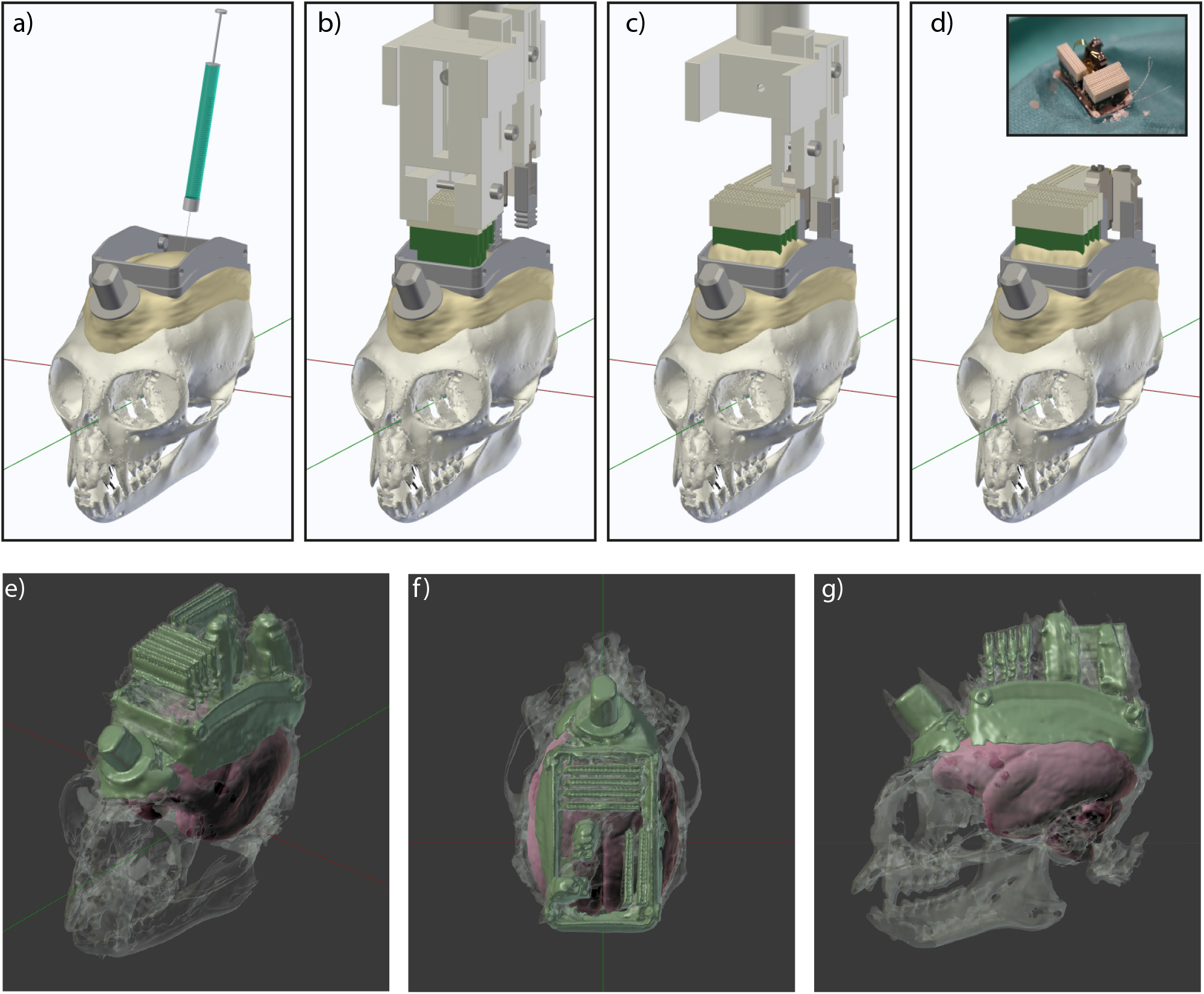
Surgery 2: Injection of the viral vector and implantation of silicon probes. a) After stereotaxic alignment of the skull via the implantation holder and the chamber, a viral vector was injected into area V6. b) A custom implantation holder, carrying connector PCBs, electrodes and microdrives was lowered into the chamber. c) First, the connector PCBs were cemented in place and the respective part of the holder was removed to ensure better access and visibility. Electrodes were then lowered sequentially into the two brain areas, and the respective microdrives were cemented into position. d) After all parts were secured, the holder was completely removed. Inset shows photograph at the end of the second surgery. e) Near-isometric projection, f) top view and g) side view of the 3D segmentation form a CT scan after the second surgery in Monkey A. Radio-opaque cement and metal parts (including connectors and microdrives) show the highest contrast and are colored in green. Bone is shown in semi-transparent gray. The fitted MRI-based template brain is shown in red.

A custom 3D printed implantation holder was then lowered into the chamber (Fig. 4b). The holder was prepared prior to the surgery to hold all necessary components for the implantation: two microdrives (with silicon probes and stabilizers attached) and six connector PCBs. The three main components (connector PCBs, V1 microdrive with probes and V6 microdrive with probes) were held by separate parts of the implantation holder, enabling independent movement in the z-axis. This independence allowed sequential implantation of the components. To this end, the holder was initially prepared such that the connector PCBs were at the lowest position and were thus implanted first (Fig. 4b). Connector PCBs were positioned via the micromanipulator just above the cement layer on the skull, and were then cemented in place. After curing, the part of the implantation holder securing the connector PCBs was removed (Fig. 4c). This resulted in better visibility and allowed for independent movement of the microdrives holding the silicon probes (Fig. 4c). Next, the probe array for area V6 was implanted. In order to insert the silicon probe into the cortex at the optimal position relative to the durotomy and the local cortical vasculature, the anterio-posterior and medio-lateral positions of the implantation holder were fine-tuned before probe insertion. After the probe was slowly inserted into the superficial part of the cortex (<500 µm), the microdrive with its attached stabilizer were cemented into the chamber. Subsequently, the part of the implantation holder that was securing the V6 microdrive was removed, too. The same procedure was performed for area V1, and the implantation holder was completely removed (Fig. 4d). Both craniotomies were then sealed with soft silicone gel (Fig. 1c).

Animals recovered very quickly after the second surgery and were brought into the recording setup within a few days. To visually inspect the position of the microdrives and PCBs, we obtained a CT scan from Monkey A after the second surgery (Figures 4 e-g). The high contrast metal parts of the connectors and microdrives with stabilizers are visible in green color. Bone is shown in semi-transparent gray and the fitted MRI-based template brain in red.

### Simultaneous recording in areas V1 and V6

After slowly lowering the probes into the brain, clear spiking activity was visible across several recording sites in areas V1 and V6 (Fig 5a).

**Figure 5.**
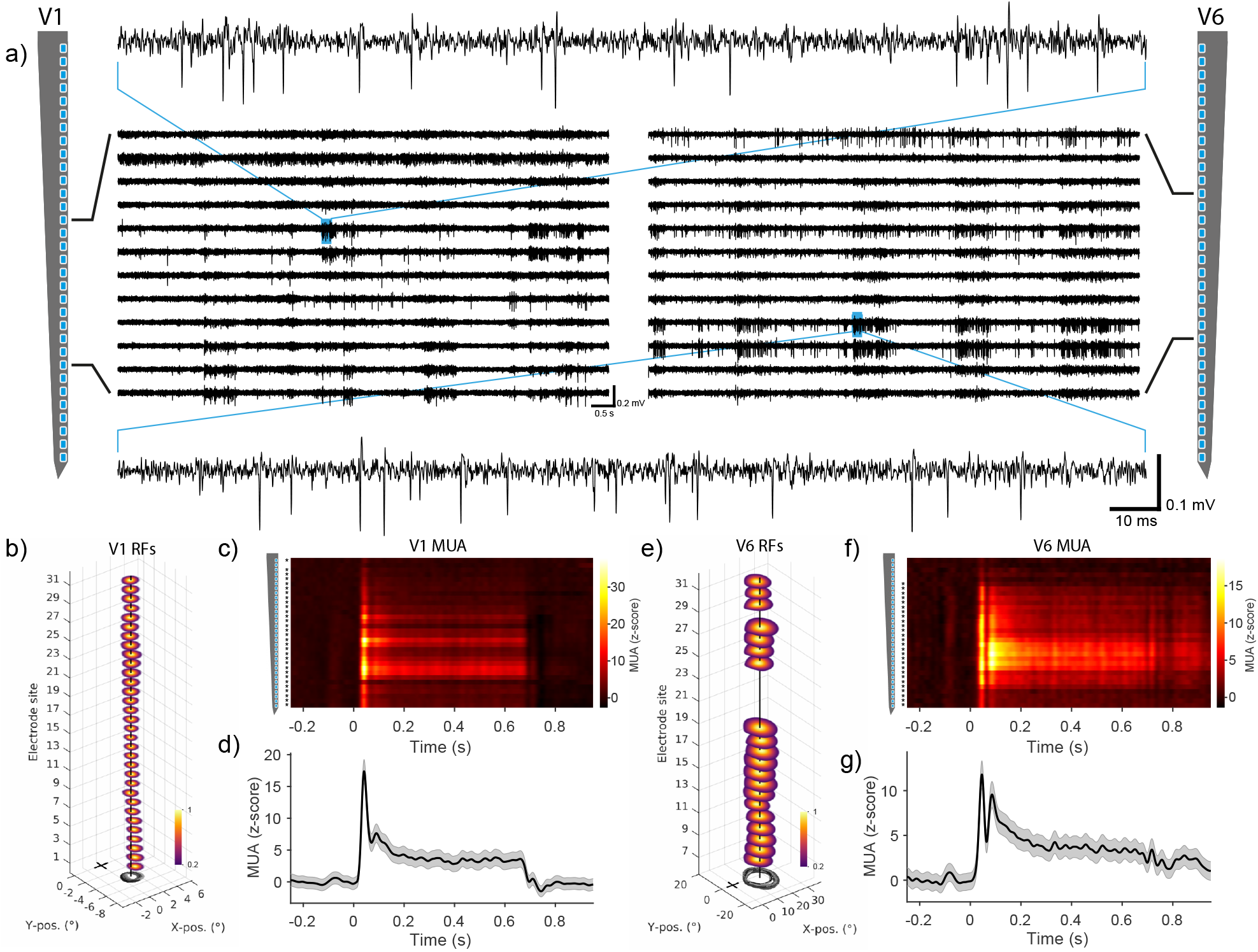
Neural recordings in areas V1 and V6. a) Band-pass filtered signal (0.3-6 kHz) from example recording sites across one shank in area V1 (left) and area V6 (right). Top and bottom traces show magnified view of the respective example signals in V1 and V6. b) Receptive field (RF) locations calculated from the normalized multi-unit-activity (MUA) of all significantly modulated sites along the example shank (n = 32 out of 32 sites). Outlines of RFs are shown at the bottom in black to gray lines from most superficial to the deepest channel. The vertical black line indicates the median RF location across all sites. The black cross marks the position of the fixation point at the center of the monitor. c) Trial-averaged MUA along the example shank around the time of visual stimulation with gratings. Asterisks on the left indicate significant modulation between pre-stimulation baseline (−0.25 to 0 s) and post-stimulus time (0 to 0.65 s) (p<0.05; Kolmogorov Smirnov test for channels with MUA>3σ). d) Average MUA ±SEM across all significantly modulated sites from the example V1 shank (n = 31 out of 32 sites). e-g) Same as b-d but for example shank in area V6 (n = 20 out of 32 sites were modulated during RF mapping; n = 27 out of 32 sites were modulated during visual stimulation with gratings). MUA was smoothed with a Gaussian window (σ = 8ms). Note different axis scaling between panel b and e. Data for RF mapping and visual stimulation with gratings were recorded in separate sessions in Monkey A.

In order to test visual responsiveness and spatial selectivity, we performed receptive field (RF) mapping with multi-unit-activity (MUA). Flashing annulus and wedge stimuli were presented while the animal was maintaining its gaze on a central fixation point. Reverse correlation analysis was used to locate RF centers across the whole monitor. A detailed account of the RF mapping procedure can be found in Jendritza et al., 2021. As expected from the implantation target position, RFs in area V1 were located in the lower right visual field (Fig. 5b). Furthermore, RFs showed substantial overlap for all electrodes along a given probe shank (Fig. 5b, black outlines at bottom).

Next, we presented static square-wave gratings to the animals. MUA following visual stimulation with gratings was visible across several recording sites and peaked shortly after stimulus onset (Fig. 5c, d). Channels were considered to contain visually modulated MUA if they fulfilled both of the following criteria: (1) The absolute magnitude of trial-averaged MUA exceeded the value of 3 STDs over the baseline (|z-score|>3) and (2) the distribution of MUA values were significantly different between baseline and stimulus period (p<0.05, Kolmogorov Smirnov test). Figure 5d illustrates the MUA averaged over all modulated sites from an example shank in V1 (n = 31 out of 32 sites). Similarly to area V1, many sites in area V6 also showed a significant spatially selective modulation (Fig 5e) (n = 20 out of 32 sites). RFs along the shank mostly overlapped, and many sites exhibited a significant MUA response after visual stimulation with gratings (Fig. 5f, g; n = 27 out of 32 sites).

### Single unit responses

Having established the overall responsiveness and visual selectivity of MUA, we next sorted spiking data into single units. Spike sorting was performed semi-automatically with the “Kilosort” algorithm (Pachitariu et al., 2016). Figure 6 depicts, in the left panel of each column, the average waveform across all 32 channels of the relevant electrode shank. Due to the fine inter-electrode spacing (25 µm), spike waveforms of each identified neuron were detectable as a spatial (and temporal) pattern across multiple sites. Raster plots and corresponding peristimulus time histograms (PSTHs) around the time of visual stimulation (black bar on top, 0.65 s duration) can be seen in the first and second row of Figure 6. The inset in the second row shows orientation tuning curves calculated from the average spiking activity during the stimulus period (0-0.65 s). Peak-normalized auto-correlograms for all spikes during the recording session are shown in the third row. The bottom row shows each unit’s firing rate over the course of a recording session, documenting that all units were stable throughout the session.

**Figure 6.**
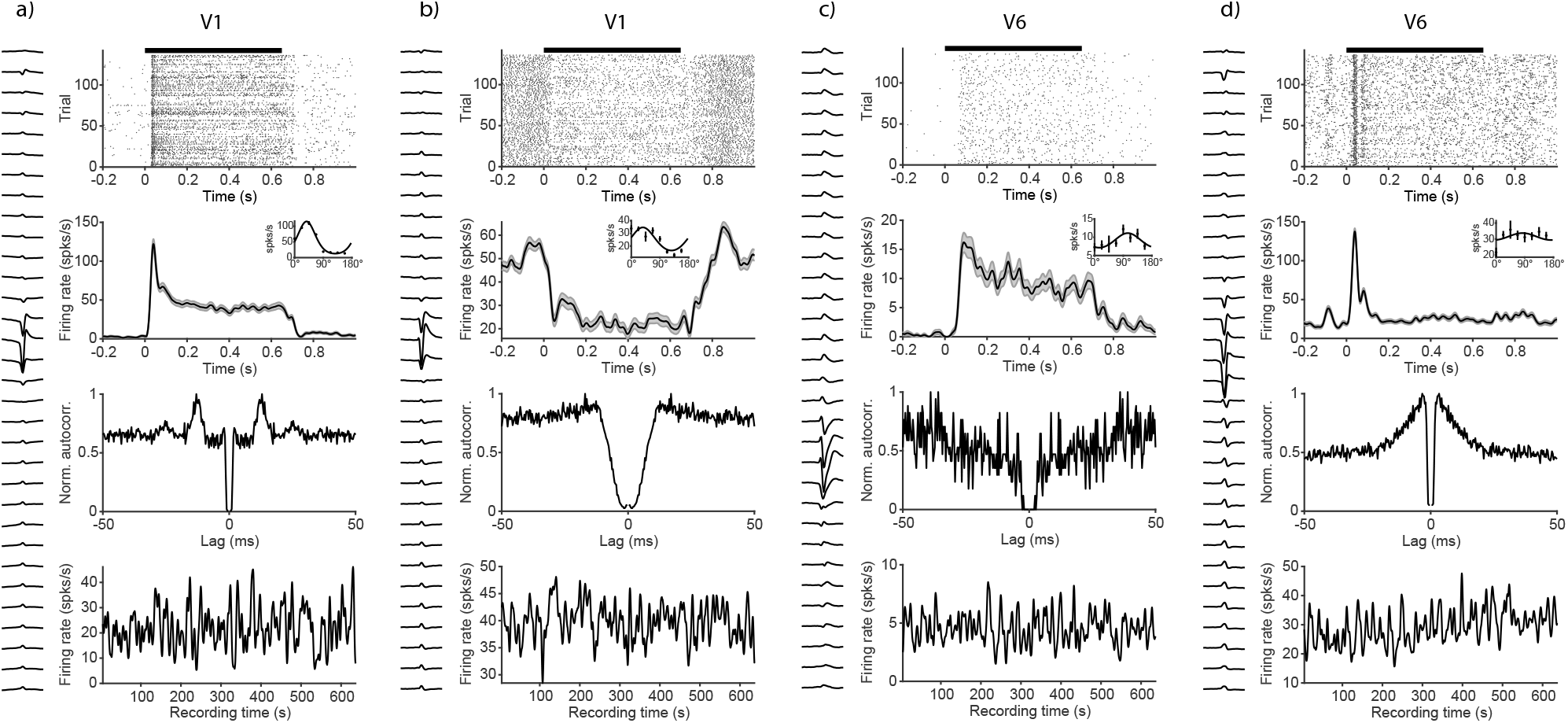
Single unit examples from areas V1 and V6. Examples of four visually modulated single units (a-d). The left side of each column shows the mean waveform across all 32 recordings sites of the electrode shank on which the largest absolute amplitude was detected. Top row: Spiking raster plot around the time of visual stimulation. Black bar on top indicates stimulus duration (0.65 s). Second row: trial averaged and smoothed (Gaussian window, σ = 10 ms) peristimulus time histogram (PSTH). Inset shows orientation tuning curves calculated from the mean activity during the stimulus period (0-0.65 s). Error bars and shaded area indicate SEM. Third row: peak-normalized auto-correlogram for all spikes across the recording. Bottom row: Smoothed firing rate (Gaussian window, σ = 2 s) across the entire session, indicating stability of the recordings. All examples from one recording session in Monkey A.

The observed single units exhibited different response characteristics, as expected from neural recordings in visual cortex. Examples in Figure 6 were selected in order to depict the variety of response profiles present in the data. The units in Fig. 6a and b were recorded in area V1. Unit a) was strongly visually driven, showed a sharp peak after stimulus onset and exhibited clear orientation tuning, reminiscent of the principal cells in V1 of the anesthetized marmoset (Yu and Rosa, 2014). The unit in Fig. 6b was suppressed during the time of visual stimulation, had a relatively high baseline firing rate, and was orientation tuned. Unit c) and d) are examples recorded in area V6. Unit c) showed a sustained activation and orientation tuning, similar to previous reports in V6 (Lui et al., 2006). In contrast, unit d) responded only transiently and exhibited only weak orientation tuning, potentially due to a non-optimal spatial frequency of the visual stimulus.

### Optogenetic stimulation of area V6

Optogenetics has become an essential tool in systems neuroscience (Deisseroth, 2015). To demonstrate that our recording approach is compatible with optogenetic stimulation techniques, we injected an adeno-associated viral vector (AAV), expressing the fast channelrhodopsin variant ‘Chronos’ (Klapoetke et al., 2014) under control of the CamKIIα promotor into area V6. Expression under the CamKIIα promotor is almost exclusively restricted to excitatory neurons (Gerits et al., 2015; Han et al., 2009; Watakabe et al., 2015). After several weeks of expression, we placed an optic fiber above the V6 craniotomy to stimulate neurons underneath the transparent silicone gel (Fig. 1c). The optic fiber was coupled to a laser that could be directly modulated with arbitrary waveforms. Stimulation was performed with sinusoidal waveforms at a peak amplitude of 25 mW.

One example trial in which stimulation was performed with an 80 Hz sinusoidal waveform is depicted in Fig. 7a. Optogenetically-induced spiking was visible across several channels. Analysis of the trial-averaged MUA revealed clear optogenetic activation time-locked to the laser waveform, for all 32 channels along the example shank (Fig. 7b). The z-scored MUA averaged across all trials and all modulated channels is presented in Fig 7c (p<0.05, Kolmogorov Smirnov test for channels with MUA>3σ, n = 32 out of 32 channels).

**Figure 7.**
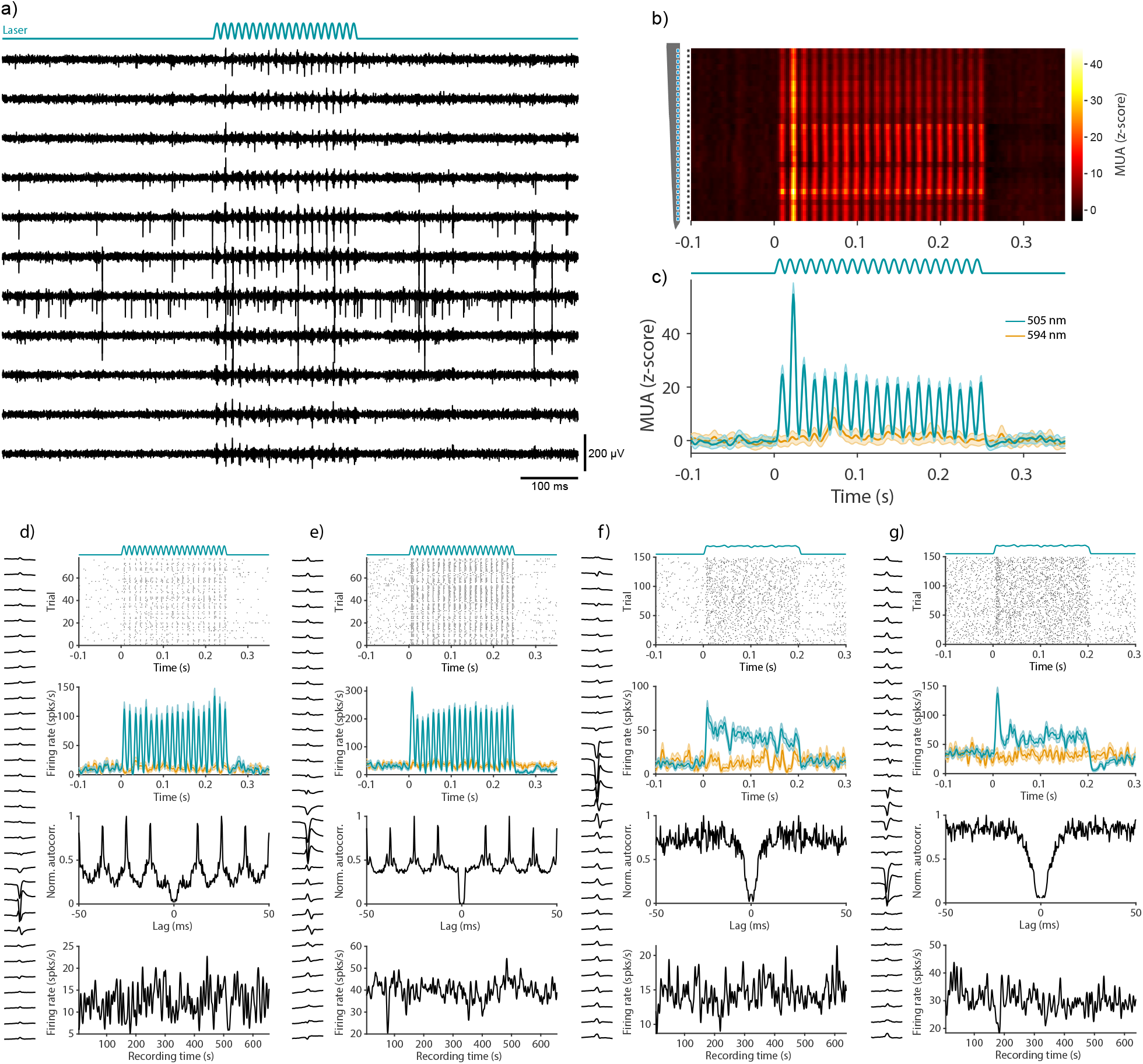
Optogenetic activation of neurons in the awake marmoset. **a)** Example traces of band-pass filtered data during optogenetic stimulation with an 80 Hz sinusoidal pattern of 250 ms duration (25 mW peak). b) Trial averaged, z-scored MUA of all recordings sites from an example shank for the 505 nm stimulation condition. Asterisks on the left indicate significant modulation between pre-stimulation baseline (−0.25 to 0 s) and stimulus time (0 to 0.25 s) across all conditions (p<0.05, Kolmogorov Smirnov test for channels with MUA>3σ) c) Average MUA across all significantly modulated channels (n = 32 out of 32 channels), for stimulation with 505 nm and 594 nm, respectively as indicated by the color legend. d-g) Four examples of optogenetically modulated single units: d) and e) from Monkey A, f) and g) from Monkey D. The left side of each column shows the mean waveform across all 32 recordings sites of the relevant electrode shank. Top row: raster plot of spikes around the time of stimulation with 505 nm. Average laser waveform across all trials is shown on top. Second row: trial averaged and smoothed (Gaussian window, σ = 2 ms) peristimulus time histogram (PSTH) for stimulation conditions with 505 nm and 594 nm (same color code as in c). Third row: peak-normalized auto-correlogram for all detected spikes during the recording session. Note the clear optogenetically induced rhythmicity in the autocorrelations of neurons in d) and e). Bottom row: Smoothed firing rate (Gaussian window, σ = 2 s) across the entire session, indicating stability of the recordings.

In order to exclude potential contamination from light-induced artifacts, we took several precautions and applied appropriate controls: First, the silicon probes used in this study are relatively robust against light artifacts (Chen et al., 2021). Furthermore, we avoided fast transients in light intensity by stimulating with low-frequency sine waves that do not contain energy in the spike frequency range. Data for MUA and SUA analysis in which optogenetic stimulation was performed, were high-pass filtered with a sharp frequency cutoff (Chebyshev Type II filter) and strong stop-band attenuation (200 dB) to remove any potential contamination from the low frequency laser signal (Wu et al., 2015). Additionally, we included a control condition, in which light with a wavelength of 594 nm with matched output power was used for optical stimulation. The opsin variant used in this study should not be activated by this wavelength (Klapoetke et al., 2014). These controls ruled out that the observed neural activation was caused by light artifacts or other non-specific effects such as heating.

Next, we spike sorted the data as described earlier in order to identify optogenetically modulated single units. Four example units are depicted in Figure 7 d-g (figure conventions are as in Figure 6). Figures 7d and e show examples from Monkey A, in which optogenetic stimulation was performed with an 80 Hz sinusoidal pattern. On each trial, sinusoidal waveforms started smoothly at the trough from an intensity of 0 mW with a peak amplitude of 25 mW. Single unit spikes were precisely time locked to the laser stimulation (Fig 7d, e). Consistent with the trial-averaged optogenetic responses, the resulting autocorrelation analysis of SUA showed a prominent peak at the reciprocal of the stimulation frequency (1/80 Hz = 12.5 ms). Figure 7f and g are additional examples from Monkey D, in which optogenetic stimulation was performed with sine waves of different frequencies (0, 10, 20, 30, 40, 50, 60, 70, 80 Hz) and randomized phases. To avoid artifacts from sharp transients in light intensity, onset and offset of the stimulation waveform were smoothed (see Materials and Methods for details). The resulting average laser intensity across all trials is shown on top of the raster plot. In both monkeys, spiking activity of single units was not affected by the control stimulation (yellow trace in second row, 594 nm), and the rates remained relatively stable throughout the recording session (Fig.7, lowermost row).

These results show that we successfully combined semi-chronic recordings in two areas with optogenetic stimulation of neurons in the visual cortex of the awake marmoset.

### Behavioral report of optogenetic stimulation

In order to test whether activation of excitatory neurons in area V6 could be behaviorally reported, we trained one animal (Monkey A) in a visual and optogenetic detection task (Fig.8a). The animal was required to briefly maintain fixation (100-150 ms) on a central fixation point. After this period, a background stimulus (full screen circular grating) was presented. After an additional 150-320 ms, a moving visual target with either low or high contrast was presented for 250 ms. Half of these trials were randomly paired with optogenetic stimulation (250 ms square pulse, 25 mW amplitude, same onset time as visual stimulus, see Materials and Methods for details). An additional condition was included in which optogenetic stimulation was performed in the absence of a visual target. The monkey was rewarded for making a saccade away from the fixation point within 500 ms from the onset time of visual and/or optogenetic stimulation. To prevent false alarms, 40% of all trials were ‘catch trials’, in which neither an optogenetic nor a visual target appeared. In these trials, the monkey was rewarded for maintaining fixation until the end of the trial. As control, we randomly interleaved trials with a sham stimulation condition. Sham stimulation was identical to real optogenetic stimulation (without a visual target), but the laser output was switched to a second optic fiber that was placed 2 mm outside the craniotomy. Importantly, sham trials were rewarded identical to real trials, such that the monkey would be able to benefit from any cues unspecific to the optogenetic stimulation.

**Figure 8.**
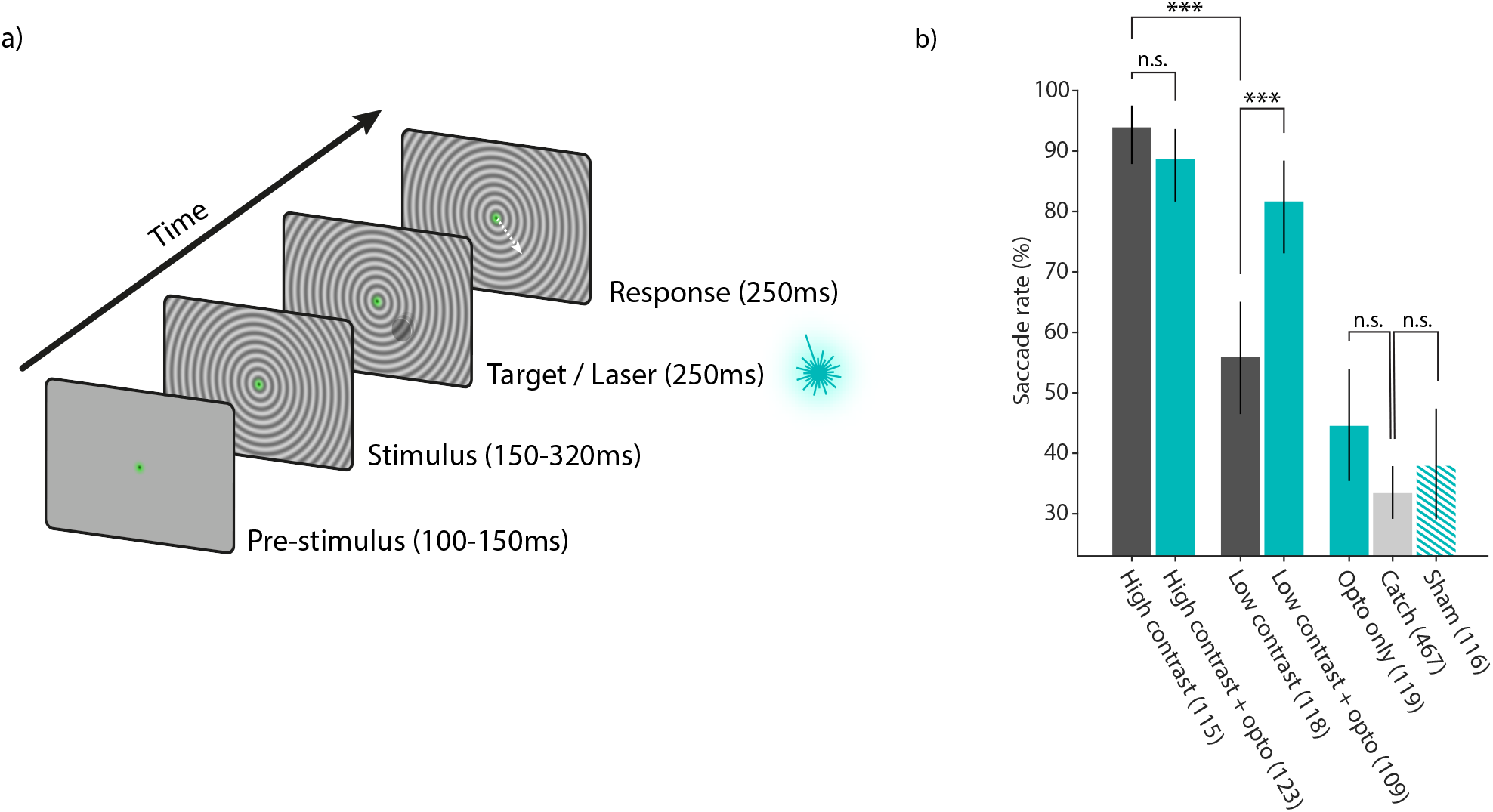
Visual and optogenetic detection task and behavioral results. a) Schematic illustration of the detection task. After a brief pre-stimulus fixation period, a background stimulus was shown, followed by the onset of a small visual target with either high or low contrast. On 50% of these trials, the visual target was paired with optogenetic stimulation. Additionally, trials without visual target were included, either with effective laser stimulation (‘Opto only’ condition) or with control laser stimulation, in which the optic fiber was placed outside the craniotomy (‘Sham’ condition), or with no laser stimulation (‘Catch’). All trial conditions except catch trials had identical timing and were rewarded if the monkey executed a saccade 50-500 ms after target or laser onset. b) Saccade rates for all task conditions. The animal showed increased detection performance (higher saccade rate) for high-contrast visual targets compared to low contrast targets (93.9% vs. 55.9%, Chi-squared test; p=3.64e-10). Pairing high-contrast visual targets with optogenetic stimulation did not result in a difference in saccade rate (Chi-squared test; p=0.283). Saccade rate increased significantly when low contrast targets were paired with optogenetic stimulation (55.9% vs. 81.7%; Chi-squared test; p=1.47e-04). Optogenetic stimulation alone was not sufficient to be detected by the animal when compared to the false alarm rate (44.5% vs. 33.4%; Chi-squared test; p=0.0521). The saccade rate in the sham stimulation control condition (laser fiber positioned 2 mm outside the craniotomy) was not different from the false alarm rate (37.9% vs. 33.4%; Chi-squared test; p=0.419). Number of trials are shown in parenthesis. Error bars indicate 95% confidence intervals.

High-contrast visual targets were correctly reported in 93.9% of trials, compared to only 55.9% in the low-contrast target condition (Chi-squared test; p=3.64e-10; n=115 high-contrast and n=123 low contrast trials). Pairing the high-contrast visual target with optogenetic stimulation did not significantly affect detection performance (Chi-squared test; p=0.283; n=115 high-contrast and n=123 high-contrast + opto trials). A different pattern was observed for low-contrast visual targets: Pairing the visual stimulus with optogenetic stimulation caused performance to improve from 55.9% to 81.7% (Chi-squared test; p=1.47e-04; n=118 low-contrast and n=109 low-contrast + opto trials). The observed increase in saccade rate indicates that the monkey was able to integrate neuronal signals from both, optogenetic and visual sources, in order to improve detection performance. The response rate to catch trials, i.e. the ‘false alarm rate’ was low (33.4%). Interestingly, optogenetic stimulation alone was no sufficient to induce a saccadic response that was significantly different from the false alarm rate (Chi-squared test; p=0.0521; n=119 opto only and n=467 catch trials). Importantly, the saccade rate in the sham control condition was not different from the false alarm rate (Chi-squared test; p=0.419; n=116 sham and n=467 catch trials).

These results show how our approach can be used to study signal integration from optogenetic stimulation during perception in the visual cortex of awake behaving marmosets.

## Discussion

Here, we demonstrate for the first time neural recordings and optogenetic stimulation in combination with behavioral manipulation in the awake behaving marmoset. Systems neuroscience relies on the constant improvement of technologies for recording and manipulation of neural circuits in vivo. Novel techniques, such as next-generation electrode technology, are therefore being developed at a rapid pace (Steinmetz et al., 2018; Hong and Lieber, 2019). Moreover, some efforts for technology development in neuroscience explicitly rely on the advantages of the marmoset model (Okano et al., 2016). Motivated by these factors, we implemented a novel approach that enables the use of modern neural probes in combination with optogenetic stimulation and behavioral manipulation in the awake, behaving marmoset. We demonstrate the functionality of our methods by obtaining multi- and single-unit recordings in two visual areas simultaneously and using optogenetic stimulation to drive neural activity and influence the animal’s behavior in a detection task.

### Advantages, drawbacks and further directions of the 3D printing-based design

Our design relies heavily on the use of 3D printing technology. 3D printing allows for rapid design adaptations, requires few mechanical constraints and enables the production of prototypes at low cost and short turnover times (Randazzo et al., 2016; Chen et al., 2017). These factors make it possible for other researchers to easily modify and improve the design presented here. There are several potential adaptations that could be useful, for example: expansion of the chamber and change in its position relative to the skull. Such modifications could enable recordings from more lateral brain areas such as area MT or IT, which are inaccessible with the current design (Suppl. Fig. 1). Moreover, the design could be adapted such that it integrates a mechanism for head fixation on the chamber (Ding et al., 2017; Johnston et al., 2018). This would make a separate headpost obsolete and thereby allow better access to frontal regions. Also, the integration of a head-fixation mechanism on the chamber might further enhance mechanical stability, which could facilitate the use with imaging techniques.

One important drawback of 3D printing methods (specifically sintering methods as used here) is that the untreated surface finish is rough. Therefore, additional steps are required if a high-precision fit (e.g. for headpost or screw threads) or a watertight sealing is necessary (Chen et al., 2017).

The weight of the complete implant, allowing recordings from 192 electrodes, amounted to approximately 8g (Fig 2c). The titanium chamber alone weighs only 1.5g and is designed to smoothly fit onto the surface of the skull with a low profile, thereby minimizing any unnecessary volume (Fig. 1a, b and 2a, b). The achieved weight minimization and the mechanical robustness of 3D printed titanium makes our design compatible with wireless recording technology. Data-loggers with batteries or wireless transmitters might be utilized, while remaining at an acceptable weight (Eliades and Wang, 2008; Roy and Wang, 2012; Walker et al., 2021). Importantly, the size and weight of implants in head-unrestrained marmosets should remain as light as possible given that these animals can perform extremely fast head movements (Pandey et al., 2020).

### Semi-chronic vs. chronic and acute recordings

Semi-chronic recording approaches, as presented here, do not require repeated insertions of electrodes into the brain for each recording session. Thus, just like chronic recordings, they can shorten the experimental preparation time and reduce the risk of infections. At the same time, such an approach retains the option of moving probes deeper into the brain after signal decay or in case the recording depth needs to be adjusted. The possibility to adapt recording depth is especially important for target locations in deeper brain structures. Thus, semi-chronic recordings with silicon probes have been recently successfully used to record neural activity from the brainstem of awake marmosets (Pomberger and Hage, 2019).

Yet, there are also advantages to other approaches such as chronic or acute recordings. In small animals, e.g. mice, immobile, chronically implanted silicon probes can provide neural recording stability over long periods of time (Okun et al., 2016; Juavinett et al., 2019; Steinmetz et al., 2021). Stability is likely related to the relative absence of movement of the mouse brain inside its skull. In marmosets, recent work has shown good recording stability with chronically implanted floating electrode (‘Utah’) arrays (Walker et al., 2021). However, long term recording stability with immobile silicon probes remains to be demonstrated. Furthermore, chronically implanted electrode arrays, such as the ‘Utah’ array do not require any movable parts and can therefore be completely sealed off after implantation, minimizing the risk of infections after surgery (Davis et al., 2020; Walker et al., 2021). Acute recording approaches on the other hand allow for repeated independent measurements and can therefore result in higher single-unit yield and make it possible to quickly change recording position (Sedaghat-Nejad et al., 2019). Thus, while semi-chronic recordings are advantageous in many circumstances, the individual experimental requirements should be considered when evaluating different recording approaches.

In this work, we performed semi-chronic recordings with silicon probe technology from passive electrodes. Yet, our design is compatible with active probes such as Neuropixels (Jun et al., 2017; Steinmetz et al., 2021) in chronic (Juavinett et al., 2019; Steinmetz et al., 2021) or semi-chronic (Vöröslakos et al., 2021) configuration. Currently, electrode shanks and microdrive-mountable components of passive silicone probes as used in this work are still smaller than those of Neuropixels probes (shank width: 25-50 µm vs. 70 µm for Neuropixels; shank thickness: 15µm vs 20µm for Neuropixels). However, active probes with fully integrated electronics and miniaturized head stages would allow for even higher channel-count recordings and will be an important next step for the advancement of neural recordings in awake marmosets.

### Optogenetic manipulation of detection behavior

We demonstrated the utility of our design by behavioral manipulation via optogenetic stimulation of area V6 in the context of a detection task. Previous work in macaques has demonstrated that optogenetic stimulation of the primary visual cortex can be readily reported via saccades (Jazayeri et al., 2012; Ju et al., 2018). These findings are consistent with the view that animals perceived phosphenes that were induced by optogenetic excitation of neurons in V1. In contrast, our own results from area V6 indicate that optogenetic stimulation alone was not sufficient to significantly modulate saccade rates. However, a clear behavioral effect was observed when laser stimulation was paired with a low contrast visual stimulus. It is known from microstimulation experiments in macaque V1 that detection sensitivity can substantially increase with behavioral training (Ni and Maunsell, 2010). Furthermore, the detection of microstimulation outside of primary sensory areas can require extended training (Histed et al., 2013). Similar changes in sensitivity thresholds have been reported for optogenetic stimulation in the somatosensory cortex (May et al., 2014). Therefore, it is plausible that further behavioral training in the marmoset would also lead to a report of optogenetic stimulation alone. This aspect should be investigated in future work.

## Materials and Methods

All animal experiments were approved by the responsible government office (Regierungspräsidium Darmstadt) in accordance with the German law for the protection of animals and the “European Union’s Directive 2010/63/EU”.

### Animals

Five adult male marmosets were implanted with chamber, headpost and reference wires. Three of these animals were subsequently injected with a viral vector in area V6, and implanted with electrodes in areas V1 and V6. The decision to use male animals was due to availability and was not part of the experimental design. Table 1 lists relevant details, procedures and outcomes for each animal.

### Stimulus presentation

Stimulus presentation was controlled by the custom-developed ARCADE toolbox (https://github.com/esi-neuroscience/ARCADE), based on MATLAB (Mathworks, USA) and C++. Stimuli were displayed on a TFT monitor (Samsung SyncMaster 2233RZ) at a refresh rate of 120 Hz. Animals were placed at a distance of 45 cm to the monitor in a dimly lit recording booth. A photodiode was placed in the top left corner of the monitor in order to determine exact stimulus-onset times.

### Eye tracking

The left eye of the animals was tracked under external infrared light illumination with a sampling rate of 1 kHz (Eyelink 1000, SR research, Canada). A 25 mm/F1.4 lens was used at a distance of 28 cm to the animal’s eye.

### Implant design and 3D printing

Designs were developed in Blender (www.blender.org), OnShape (https://www.onshape.com/), and Solidworks (https://www.solidworks.com/). 3D renderings were generated in Blender. The skull template shown in Figures 1 and 2 was segmented with 3D Slicer (https://www.slicer.org/) based on high-resolution CT data from a marmoset skull archived on the MorphoSource data base (https://doi.org/10.17602/M2/M5203/). Chambers and microdrive stabilizers were printed via direct metal laser sintering from grade 5 (Ti6Al4V) titanium (Materialise, Belgium). Microdrives were glued to the stabilizers with cyanoacrylate glue. Lids were printed via selective laser sintering from PA12 nylon (Shapeways, USA). To ensure watertight sealing, a thin layer of silicone (Kwik-Sil, World Precision Instruments, USA) was applied to the small ridge inside the lid that served as contact area between chamber and lid. All custom implantation holders and guides were printed from standard resins via stereolithography on a “Form 1” printer (Formlabs Inc., USA). Design files for 3D printing can be found at https://github.com/PJendritza/Marmo/.

### Anesthesia

Anesthesia for all surgeries was induced with an intramuscular (i.m.) injection of a mixture of alfaxalone (8.75 mg/kg) and diazepam (0.625 mg/kg). Tramadol (1.5 mg/kg) and metamizol (80 mg/kg) were injected i.m. for initial analgesic coverage. Subsequently, a continuous intravenous (i.v.) infusion was provided through the lateral tail vein. The i.v. mixture contained glucose, amino acids (Aminomix 1 Novum, Fresenius Kabi, Germany), dexamethasone (0.2-0.4 mg·kg-1·h-1), tramadol (0.5-1.0 mg·kg-1·h-1) and metamizol (20-40 mg·kg-1·h-1). The maximal infusion rate was 5 ml·kg-1·h-1. Animals were breathing spontaneously throughout the surgery via a custom 3D printed face mask that applied isoflurane (0.5-2% in 100% oxygen). Heart rate, respiration rate and body temperature were constantly monitored (Model 1030 Monitoring Gating System, SAII, USA).

### Implantation of chamber and headpost

After placing the animal in a stereotaxic apparatus for the first surgery, an incision was made on the dorsal part of the skull. The temporal muscle was slightly retracted (<5 mm from the superior temporal lines) and all soft tissue was completely removed from the bone surface. The bone was first cleaned by mechanical abrasion, then scrubbed with 5% H_2_O_2_ and rinsed with saline. For an optimal bonding between cement and bone, the skull surface was roughened with a metal brush, and any remaining dust was removed. After the bone was completely clean and dry, we applied a thin layer of light-curable dental adhesive (All-Bond Universal, BISCO). After drying and curing with blue light, we applied a thin layer (<1 mm) of dental cement on top of the adhesive. Once the cement was cured, a small bur hole was drilled just anterior of the chamber. Two platinum wires (PT-5T, Science Products) were implanted epidurally at this location and served as backup reference wires for the recordings (the actual reference wires were later implanted subdurally in the second surgery).

### Injection of the viral vector

Viral vectors (AAV1.CamKIIa.Chronos-eYFP-WPRE) were injected with a microinjector pump (UMP3-1, World Precision Instruments), holding a 10uL microsyringe (NanoFil syringe, World Precision Instruments) to which a 35G injection needle was attached. A durotomy of approx. 1.5 mm was performed with a bent 25G cannula, and the vector was injected at two depths (−1.4 mm and -0.5 mm from the surface). A volume of 2.5 µl at each depth was injected at a speed of 200 nL/min (total injected volume = 5.0 µl). To ensure sufficient diffusion of the viral vector, we waited 10 min after the each injection before moving or retracting the needle.

### Silicon probes

Silicon probes were semi-chronically implanted in areas V1 and V6, mounted on one microdrive per area (Nano-Drive CN-01 V1, Cambridge NeuroTech, UK). Two 32-channel shanks with 250 µm spacing were implanted in V1, and four 32-channel shanks in V6 (H2 probe, Cambridge NeuroTech, UK). Electrode implantation was performed directly following the injection of the viral vector. Electrode tips were disinfected shortly before the implantation by dipping them twice in 70% ethanol for 45 s. After the electrodes were in place and the cement was hardened, craniotomies were sealed by applying several drops of soft silicone gel (DOWSIL 3-4680, Dow Corning).

### Acquisition and processing of neural data

Neural signals were recorded through active, unity gain head stages (ZC32, Tucker Davis Technologies, USA), digitized at 24,414.0625 Hz (PZ2 preamplifier, Tucker Davis Technologies, USA) and re-sampled offline to 25 kHz. Sample-by-sample re-referencing was applied by calculating the median across all channels for each shank and subtracting this signal from each channel of the corresponding shank (Jun et al., 2017). Data was band-pass filtered for spiking activity either with a 4th-order Butterworth filter (0.3-6 kHz) or, in case optogenetic stimulation was performed, with a 40th-order Chebyshev Type II filter (0.3-8 kHz) with a stop-band attenuation of 200 dB to exclude any contamination from lower frequencies. For further analysis, multi-unit activity (MUA) was calculated by full-wave rectification, filtering with a 6th-order low-pass Chebyshev Type II filter (stopband edge frequency of 500 Hz, stopband attenuation of 50 dB) and down-sampling to 1 kHz.

### Optogenetic stimulation

Optogenetic stimulation was performed with a laser beam combiner (LightHUB, Omicron laserage), housing a 100 mW diode laser with a wavelength of 505 nm (LuxXplus 505-100) with direct modulation and a 100 mW DPSS laser with a wavelength of 594 nm (OBIS 594-100) with direct modulation. The combined lasers were coupled to a 50μm/0.22NA optic fiber which was connected to a fiber optic cannula (200 µm core diameter, 0.39 NA, Doric Lenses Inc.). The cannula was held by a micromanipulator (SM-25C, Narishige) and was positioned approx. 4 mm above the craniotomy during recording/stimulation sessions. Laser power was calibrated prior to the experiments with a photodiode-based optical power meter (PM130D, Thorlabs). Output power was measured at the tip of the fiber optic cannula. Laser waveforms were generated by a real-time signal processor (RZ2 bioamp processor, Tucker Davis Technologies, USA). To avoid artifacts arising from sharp transients in laser intensity (Cardin et al., 2010), we only used smooth on and offsets (Wu et al., 2015). This was done by using one half of a sine wave as a taper at the beginning and end of any sharp signal (5 ms trough-to-peak time, with the trough having an intensity of 0 mW).

### CT scans and segmentation

CT scans were performed under brief anesthesia induced with an intramuscular (i.m.) injection of a mixture of alfaxalone (8.75 mg/kg) and diazepam (0.625 mg/kg). The head of the animal was stabilized via the headpost for the duration of the scan. CTs were performed with a Planmeca

ProMax 3D Mid scanner (Planmeca Oy, Finland) at 90 kV and 10 mA with a voxel size of 150 μm (isotropic). Segmentation of CT data was performed with 3D Slicer. Models were exported as STL files and imported into Blender for alignment.

### Spike sorting and single unit analysis

Spike sorting was performed offline with Kilosort (Pachitariu et al., 2016). Average spike waveforms were calculated from the trimmed mean (5% outlier exclusion). Autocorrelation functions were generated at a resolution of 0.33 ms and normalized by dividing by the maximum value after removal of the central peak.

### Receptive field mapping

All details about the RF mapping procedure have been described previously in Jendritza et al., 2021. RF mapping was performed with stimuli consisting of black wedges and annuli of various orientations and sizes, presented on a gray background for a duration of eight frames (120 Hz monitor refresh rate). For RF calculation, MUA data was cut into epochs of 280 ms (from 100 ms before to 180 ms after stimulus onset). Epochs were included in the analysis if the eye position remained inside the fixation window throughout the epoch. For noise-rejection purposes, we excluded epochs in which the standard deviation of MUA across time was more than 10-times larger than the median standard deviation across all epochs. Sites were considered to be modulated if the mean MUA from at least three different wedge stimuli and at least three different annulus stimuli evoked a response that was significantly larger (paired t-test, alpha = 0.01) than the MUA during baseline (100 ms to 0 ms prior to stimulus onset). For plotting, MUA was normalized per site to have a value between zero and one. RF plots and outlines were generated by truncating the normalized MUA at a value of 0.2.

### Passive fixation task

A passive fixation task was used to measure neural responses following visual stimulation with gratings. At the beginning of each trial, the animal was required to maintain its gaze at a central fixation point within a window of 1.4° radius for 100-120 ms. After this period, a static square-wave grating was presented for 650 ms at a Michelson contrast of 80%. The size and orientation of the grating was selected at random for each trial. Possible values for the grating radius (in degrees of visual angle) were: 5°, 7.25°, 9.5°, 11.75° and 14°. Possible values for the grating orientation were: 22.5°, 45°, 67.5°, 90°, 112.5°, 135°, 157.5° and 180°. After stimulus offset, the animal was required to maintain it gaze in the fixation widow for another 100 ms. After a correct trial, a picture of a marmoset face was displayed in the center of the monitor, and the animal was rewarded. The amount of reward was 0.07 ml per trial at the start of the session and increased by 0.02 ml for every 10 ml consumed (capped at 0.1 ml per trial). Reward was provided via a lick spout and consisted of diluted *gum arabic*.

### Visual and optogenetic detection task

At the beginning of each trial of the detection task, the animal was required to position its gaze at a central fixation point within a window of 1.5° radius for 100-150ms. After this period, a background stimulus was presented, while the monkey maintained fixation. The background stimulus was a full-screen circular grating, concentric to the fixation point and either contracting towards or expanding from the fixation point, each in a random half of the trials (contrast = 40%, spatial freq. = 2 cycles/°, temporal freq. = 1 cycle/s). At 150-320 ms after the onset of the background stimulus, a black, moving circular patch (1.8° diameter, moving at 5.74 °/s, linear motion, random direction) with either high (50%) contrast or low contrast (7.8%) was presented for 250 ms. The center of the movement path of the circular patch was fixed in the lower right quadrant, where the receptive fields of the optogenetically responsive V6 cells were located. Additionally, a condition was included in which only optogenetic stimulation was performed in the absence of a visual target. Furthermore, a control “sham” stimulation condition was included, with sham trials being identical to real optogenetic stimulation trials (without visual target), but with the laser output switched to a second optic fiber that was placed 2 mm outside the craniotomy. All of these “go” trials (60% of all trials) were categorized as hits if the animal made a saccade away from the fixation point within 500 ms after the onset of the moving circular patch or the laser. Responses that were faster than 50 ms were categorized as early responses and were not rewarded. 50% of trials with a visible target were coupled with optogenetic stimulation that consisted of a 250 ms square pulse with an amplitude of 25 mW. The onset timing for visual and optogenetic stimulation was determined by the computer controlling the visual stimulation. We did not compensate for any delay between trigger onset and actual onset of the visual stimulus on the monitor. In the remaining “catch” trials (40% of all trials), no visual or optogenetic target was presented, and the monkey was rewarded for maintaining its gaze at the fixation point for 800 ms. After a correct saccade, or a correct rejection (maintained fixation), a picture of a marmoset face was displayed in the center of the monitor, and the animal was rewarded. The amount of reward was 0.0625 ml per trial at the start of the session and increased by 0.02 ml for every 10 ml consumed (capped at 0.1ml per trial).

In the detection task described above, catch trials were longer than the average go trial. Thus, simply calculating saccade rates from catch trials would lead to an overestimation of the true false alarm rate, because the monkey had more time to perform a saccade in a catch trial than in a go trial. False-alarm rate calculation was therefore performed in the following way: One randomly selected catch trial with false alarm was compared with the timing of a randomly selected go-trial. If the time of the false alarm from the selected catch trial fell within the time window in which the monkey would have performed a hit, the trial was categorized as a false alarm. If the false alarm timing was such that the monkey would have missed the target, the trial was categorized as correct rejection. This random pairing was performed for n=467 random pairs of trials, as this was the expected number of catch trials (40% of all trials), given the total number of hits and misses performed by the animal (n=700). The proportion of false alarms and the respective binomial confidence intervals were then calculated for this random sample. This procedure was repeated 1000 times, and the false-alarm rates and confidence intervals from all shuffling iterations were averaged.

## Acknowledgements

We thank Marianne Hartmann for her constant support in training the animals, Martin Vinck for providing access to his GPU systems and Gustavo Rohenkohl for his feedback on the manuscript.

This work was supported by DFG (SPP 1665 FR2557/1-1, FOR 1847 FR2557/2-1, FR2557/5-1-CORNET, FR2557/6-1-NeuroTMR, FR2557/7-1-DualStreams to P.F.), EU (HEALTH F2 2008 200728-BrainSynch, FP7-604102-HBP, FP7-600730-Magnetrodes to P.F.), a European Young Investigator Award to P.F., National Institutes of Health (1U54MH091657-WU-Minn-Consortium-HCP to P.F.), the LOEWE program (NeFF to P.F.).

## Supplementary Figures

**Supplementary Figure 1.**
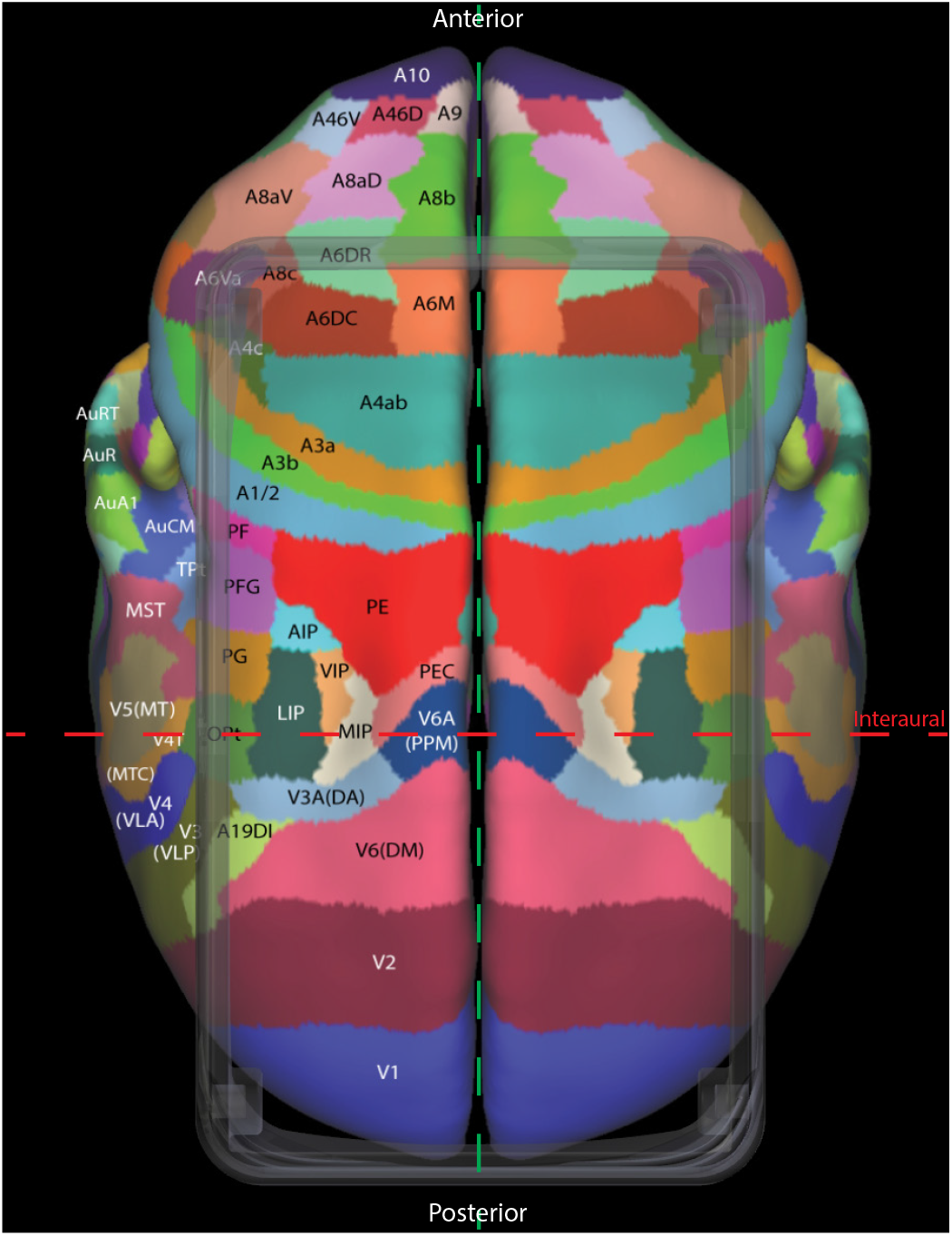
Accessible cortical areas. Top view of the chamber and cortical brain areas directly underneath. Red dashed line indicates interaural axis. Green dashed line indicates anterior-posterior axis. Area segmentation and labels from Paxinos et al. (2012).

**Supplementary Figure 2.**
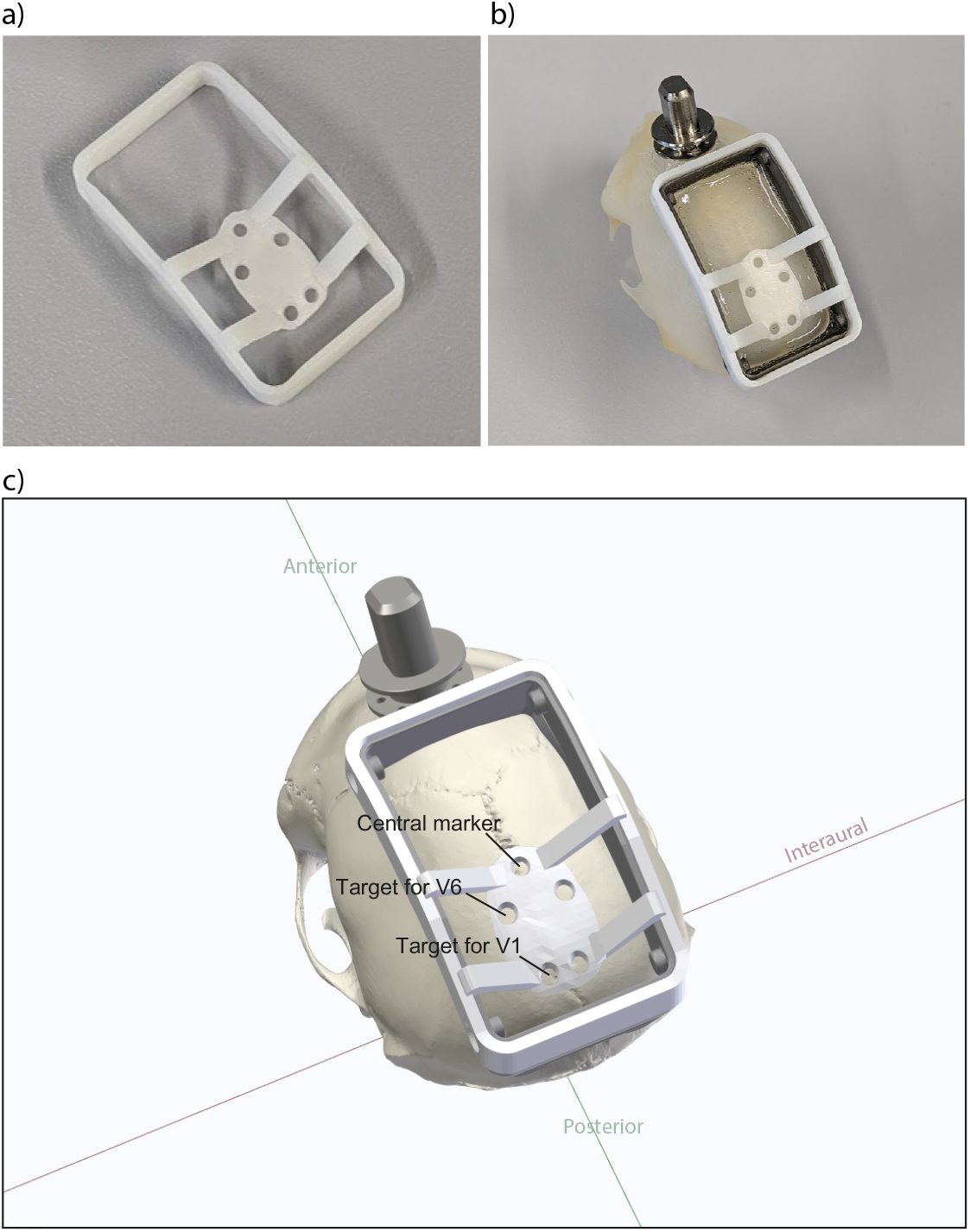
3D printed implantation target guide. Photograph of the implantation guide (a) before and (b) after placement on the chamber. c) 3D rendering of the implantation guide placed on the chamber. The guide hole for the central marker indicates the anterior-posterior and medio-lateral center of the stereotaxic coordinate system. Guide holes for areas V1 and V6 are indicated for the left hemisphere.

